# AMP-Diffusion: Integrating Latent Diffusion with Protein Language Models for Antimicrobial Peptide Generation

**DOI:** 10.1101/2024.03.03.583201

**Authors:** Tianlai Chen, Pranay Vure, Rishab Pulugurta, Pranam Chatterjee

## Abstract

Denoising Diffusion Probabilistic Models (DDPMs) have emerged as a potent class of generative models, demonstrating exemplary performance across diverse AI domains such as computer vision and natural language processing. In the realm of protein design, while there have been advances in structure-based, graph-based, and discrete sequence-based diffusion, the exploration of continuous latent space diffusion within protein language models (pLMs) remains nascent. In this work, we introduce AMP-Diffusion, a latent space diffusion model tailored for antimicrobial peptide (AMP) design, harnessing the capabilities of the state-of-the-art pLM, ESM-2, to *de novo* generate functional AMPs for downstream experimental application. Our evaluations reveal that peptides generated by AMP-Diffusion align closely in both pseudo-perplexity and amino acid diversity when benchmarked against experimentally-validated AMPs, and further exhibit relevant physicochemical properties similar to these naturally-occurring sequences. Overall, these findings underscore the biological plausibility of our generated sequences and pave the way for their empirical validation. In total, our framework motivates future exploration of pLM-based diffusion models for peptide and protein design.

## 1 Introduction

In the evolving landscape of therapeutics, proteins and peptides have carved a niche, offering targeted solutions for a myriad of disease indications (1). Among these, antimicrobial peptides (AMPs) present a compelling alternative to traditional antibiotics, especially in the face of rising drug-resistant pathogens. These short peptides, characterized by their compact and adaptable structures, have the potential to revolutionize therapeutic interventions for diseases driven by bacteria, fungi, parasites and viruses (2). However, the current paradigm for engineering these peptides is predominantly anchored in high-throughput screening and rational design, aiming to enhance *in vivo* stability, solubility, and strain specificity while mitigating aggregation (3). The inherent flexibility of peptides, while advantageous for clinical applications, complicates computational design, as traditional structure-based approaches are often unable to adeptly handle the dynamic, conformationally unstable nature of these molecules (4). In total, these features underscore the pressing need for a sequence-centric peptide generation platform, streamlining the design of peptides primed for empirical validation.

Recent advancements in generative artificial intelligence (AI) have highlighted the efficacy of diffusion models (5), particularly in domains such as computer vision and natural language processing. In the specialized field of protein design, there has been a growing interest in leveraging diffusion-based methodologies, and encompassing approaches like structure-based diffusion, sequence-based diffusion, and graph-based diffusion models, each demonstrating promising outcomes in their respective tasks. For instance, RFDiffusion (6) has been pivotal in the design of protein monomers, binders, and symmetric oligomers, integrating structural data for enhanced outcomes. On the other hand, EvoDiff (7) excels in generating intrinsically disordered regions with evolutionary sequence information. Furthermore, the graph-based diffusion approach has made significant strides in areas like antibody design (8) and protein-ligand docking (9). Concurrently, the development of protein language models (pLMs) such as ESM-2 (10), ProtT5 (11), ProGen (12), and ProtGPT2 (13), has significantly propelled our understanding and design capabilities for proteins. The evolving synergy between diffusion models and pLMs presents an exciting frontier for innovative methods in protein research. However, the integration of latent diffusion techniques with the foundational knowledge of pLMs remains a relatively unexplored territory. This fusion is poised to enrich diffusion models with prior evolutionary knowledge and offer more nuanced control within the latent space, marking a promising advancement in the computational design of proteins.

In this study, we introduce the first latent-space diffusion pLM, named AMP-Diffusion. This model uniquely harnesses the diffusion model framework to generate novel antimicrobial peptides (AMPs), serving as a demonstration of latent diffusion techniques for protein sequence generation. Leveraging latent space of ESM-2 (10), a state-of-the-art pre-trained pLM, AMP-Diffusion employs a diffusion framework that introduces Gaussian noise during the forward phase and adeptly reverses this process to reconstruct the peptide embeddings from their noised inputs. Post-training, AMP-Diffusion is capable of generating AMPs that not only exhibit low perplexity but also display a high degree of diversity and similarity, reflecting the complex nature of real AMPs.

## 2 Methods

### AMP-Diffusion

Diffusion models are designed to approximate an unknown data distribution, *p*(**x**), by creating a bridge between a simple Gaussian distribution and a complex data distribution. The forward diffusion process starts with a data sample, **x** *∼ p*(**x**), and introduces a series of latent variables {**z**_0_, **z**_1_,, *…***z**_*T*_} that transition from the data distribution towards a Gaussian distribution over increasing timesteps. This transition is governed by a Markov chain, where the variance *β*_*t*_ at each step determines the amount of Gaussian noise added. Specifically, *α*_*t*_, which is a cumulative product of the variances (1 *− β*_*i*_) up to time *t*, dictates the noise level. The equation 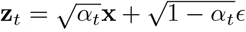 describes how the original data is perturbed, with *ϵ* being Gaussian noise. In the generative process, this forward diffusion is inverted. Starting with pure Gaussian noise, **z**_*T*_ *∼ N* (0, 1), the model iteratively denoises the noise to produce samples resembling the original data distribution with the Loss 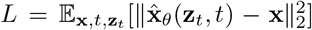. For sampling from the diffusion model, one can start with Gaussian noise in the reverse process, and then obtain a sample from the data distribution, **z**_0_, by DDPM (5), Denoising Diffusion Implicit Models (DDIM), and other samplers (14).

For latent diffusion-based generation of AMPs (Figure 1A), the ESM-2 encoder is employed to map peptide sequences into a continuous latent space represented as **x** *∈ ℝ*^*𝓁×d*^, on which Gaussian noise is added at each time step in the forward process. Simultaneously, a denoiser is trained to reconstruct the embeddings disturbed by this noise. In the reverse process, the denoiser works on Gaussian noise to revert it back into latent embeddings that align with the ESM-2 latent space. The training objective focuses on minimizing the *l*_2_ loss between the predicted and original *x*_0_. The latent representations are then decoded into peptide sequences using the ESM-2 language model head. In the model configuration (Figure 1B), the denoising process employs pre-trained ESM-2-8M attention blocks. A positional encoding is utilized for the time embedding, integrated into protein embeddings with a scaling factor and a bias adjustment (15). The output from these transformer layers is then directed into a straightforward multilayer perceptron (MLP) for further processing.

**Figure 1:**
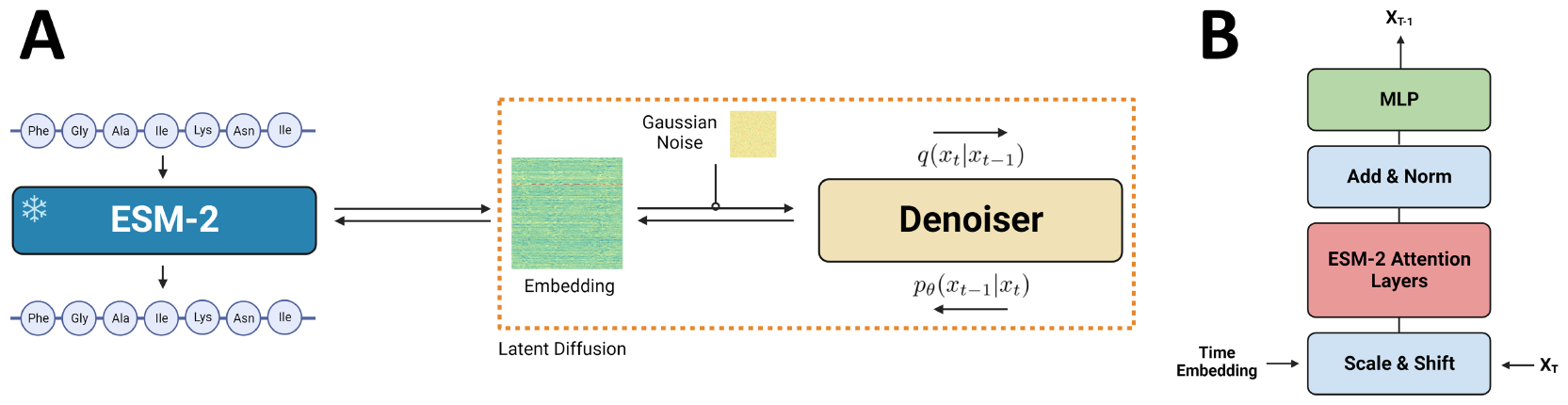
A) Overview of latent diffusion with ESM-2. AMP sequences are fed into ESM-2 for embedding generation. In forward process (*q*), Gaussian noise is introduced into these latent embeddings. In the reverse process (*p*_*θ*_), the model is trained to denoise and reconstruct latent embeddings. B) Denoiser architecture. The model consists of ESM-2 attention layers and simple multilayer perceptron (MLP) networks.

### Data

A dataset comprising 195,121 peptide sequences was collected from recognized databases, including dbAMP (16), AMP Scanner (17), and DRAMP (18). Duplicate sequences were systematically removed to ensure data integrity. All sequences selected for the training set adhere to a maximum length of 40 amino acids. Sequences not reaching the 40 amino acid length specification were supplemented with the padding token. For generation and evaluation, 50,000 sequences were sampled from AMP-Diffusion and other relevant models (HydrAMP (19), PepCVAE (20), and AMPGAN (21), see 5.2).

### ESM-2 Pseudo-Perplexity

The model’s generation quality was assessed using the ESM-2 (10) pseudo-perplexity metric. Typically, a lower pseudo-perplexity value indicates higher confidence. Specifically, the pseudo-perplexity is computed as the exponential of the negative pseudo-log-likelihood of a sequence. This metric yields a deterministic value for each sequence but necessitates L forward passes for computation, where L represents the input sequence length. It is formally defined as: 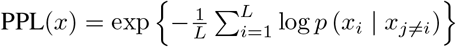

### Entropy and Diversity

To evaluate the diversity and similarity of the generated peptide sequences, two metrics are utilized: Shannon entropy and Jaccard similarity. Shannon entropy serves as a measure of the sequence’s uncertainty or randomness, reflecting its information content and complexity. It is calculated using the formula *H*(*X*) = *p*(*x*) log_2_ *p*(*x*), where *p*(*x*) denotes the probability of occurrence of each amino acid in the peptide. A higher entropy value indicates a more diverse peptide sequence, assessed by the frequency of each amino acid. On the other hand, the Jaccard similarity (JS) coefficient is employed to compare the similarity of generated peptide sets to the training data, specifically at the level of k-mers. Defined as 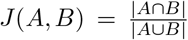, this coefficient measures the similarity between two sets, allowing for the assessment of shared and unique motifs in peptide sequences. For this study, 3-mers and 6-mers are particularly analyzed to gauge the effectiveness of the generated sequences.

### Physicochemical Properties

Generated peptides from AMP-Diffusion were benchmarked against those from other models and compared with natural AMPs (the training data). The evaluation of physicochemical properties, including charge, hydrophobicity, aromaticity, and isoelectric point (pI), was conducted using the modlAMP toolkit (22). The isoelectric point (Pi), the pH at which a peptide’s net charge is zero, was determined using specific pK values for amino acid residues. Charge calculations were performed following Bjellqvist’s method (23), which assesses the net charge under various pH conditions. Peptides with higher charge and isoelectric point values are generally more favorable for AMP function (24). Hydrophobicity was quantified using the Eisenberg scale (25), a measure of amphiphilicity in peptide structures (26). Aromaticity was evaluated based on the occurrence of phenylalanine, tryptophan, and tyrosine (27). Similarity in aromaticity and hydrophobicity metrics to real AMPs indicate the potential functional validity of the generated peptides.

### External Classifier

The HydrAMP classifier from the state-of-the-art model for AMP generation (19) serves as an external validation tool to ascertain the antimicrobial properties of generated peptides. This classifier is proficient in determining whether a peptide is antimicrobial and in estimating its minimal inhibitory concentration (MIC) against *E. coli*. The classifier consists of two distinct networks: the MAMP, which predicts the probability of a peptide being antimicrobial, expressed as *P*_amp_; and the MMIC, which calculates the likelihood of the peptide’s activity against *E. coli*, denoted as *P*_mic_. For AMP detection, the MAMP network is trained using AMP Scanner (17), utilizing a comprehensive dataset of known AMP sequences. In contrast, the MMIC network is tailored with a unique architecture to process MIC data, enabling precise predictions of peptide efficacy.

## 3 Results

Table 1 delineates the performance of AMP-Diffusion in comparison to other prevalent models, as assessed by metrics such as perplexity, entropy, and Jaccard similarity indices for 3-mers (JS-3) and 6-mers (JS-6). AMP-Diffusion is noted for its superior performance, evidenced by the lowest perplexity at 12.84 and the highest entropy at 3.17, indicative of its effective sequence generation. While all models demonstrate high similarity scores at the 3-mer level (0.99), AMP-Diffusion achieves a markedly higher Jaccard similarity at the 6-mer level (0.028), reflecting its robust capability in capturing complex sequence motifs relevant for AMP design.

**Table 1:** Evaluation of AMP-Diffusion and relevant models.

| Model | Architecture | Perplexity | Entropy | JS-3 | JS-6 | $P_{amp}$ | $P_{mic}$ |
| --- | --- | --- | --- | --- | --- | --- | --- |
| AMP-Diffusion | Diffusion | $12.84 \pm 4.53$ | $3.17 \pm 0.56$ | 0.99 | 0.028 | 0.81 | 0.50 |
| Train/Real AMPs | - | $16.23 \pm 3.80$ | $3.23 \pm 0.40$ | - | - | - | - |
| HydrAMP (19) | CVAE | $17.27 \pm 6.02$ | $2.82 \pm 0.48$ | 0.99 | 0.014 | 0.77 | 0.49 |
| PepCVAE (20) | CVAE | $19.82 \pm 3.83$ | $3.12 \pm 0.36$ | 0.99 | 0.020 | 0.41 | 0.20 |
| AMPGAN (21) | GAN | $17.70 \pm 4.08$ | $2.90 \pm 0.61$ | 0.99 | 0.016 | 0.54 | 0.32 |

In the classification assessment, peptides were evaluated for their likelihood of being antimicrobial with a threshold for *P*_amp_ set above 0.8. Additionally, peptides were assessed for their activity potential against *E. coli* with a threshold for *P*_mic_ set above 0.5. AMP-Diffusion exhibits a notable proficiency, with the highest proportion of peptides exceeding both thresholds. This performance not only suggests a higher likelihood of peptides being classified as AMPs but also indicates a greater probability of exhibiting effective activity, thus highlighting the robustness of AMP-Diffusion in generating potentially active peptides.

The physicochemical property analysis is depicted in Figure 2. The AMP-Diffusion model generates peptides with isoelectric point (pI) and charge distributions that are closely aligned with those of the training dataset, exhibiting a broad interquartile range. This suggests that AMP-Diffusion can generate a diverse array of peptides with pI values that are potentially advantageous for biological activity. In contrast, HydrAMP and AMPGAN exhibit higher pI and charge values; however, these largely fall outside the distribution of natural AMPs, which may put into question their biological relevance. PepCVAE demonstrates a performance comparable to AMP-Diffusion in terms of pI and charge. Regarding hydrophobicity and aromaticity, AMP-Diffusion and other models generally match the distribution of real AMPs, with the exception of HydrAMP. Notably, the peptides from AMP-Diffusion display a wide range of hydrophobicity and aromaticity, indicating the model’s ability to synthesize peptides that extend beyond the diversity found in natural AMPs. This expanded range could potentially lead to novel peptides with unique properties suitable for various therapeutic applications.

**Figure 2:**
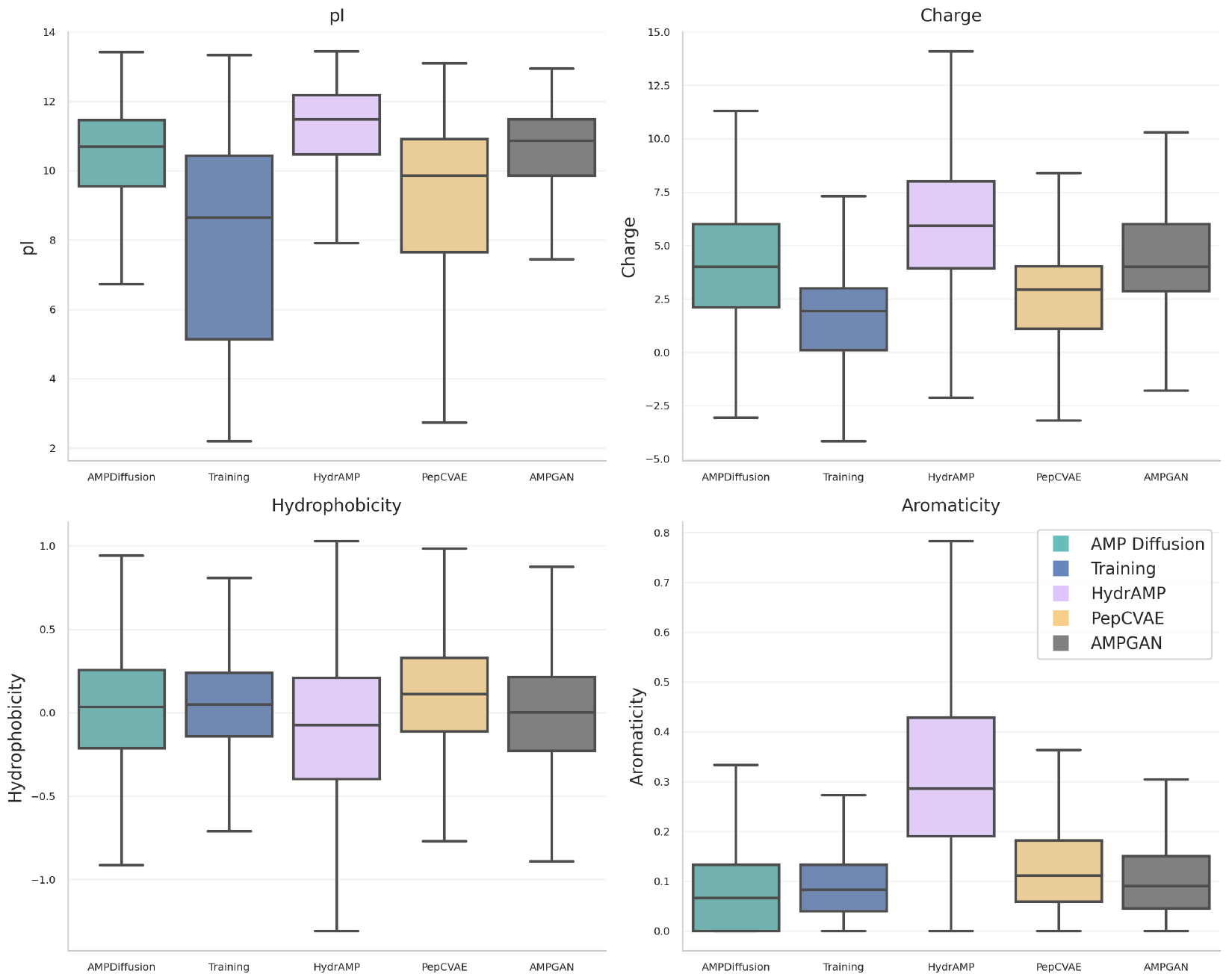
Distribution of physicochemical properties including isoelectric point (pI), charge, hydrophobic ratio, and aromaticity for real AMPs from the training dataset compared to peptides generated through unconstrained generation by AMP-Diffusion, HydrAMP, PepCVAE, and AMP-GAN. The sample size for each group is (n=50,000).

**Figure 3:**
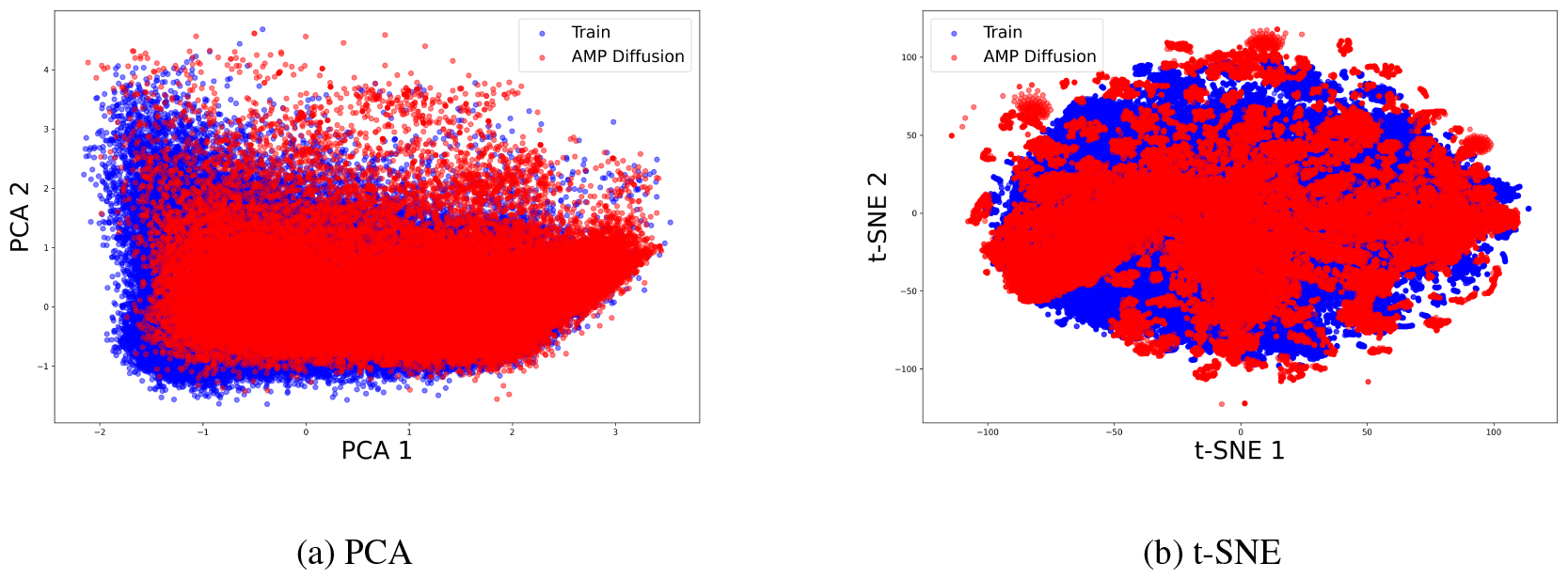
Embedding visualizations of PCA and t-SNE. Blue dots represent training data embeddings, while red dots denote embeddings generated by AMP-Diffusion.

**Figure 4:**
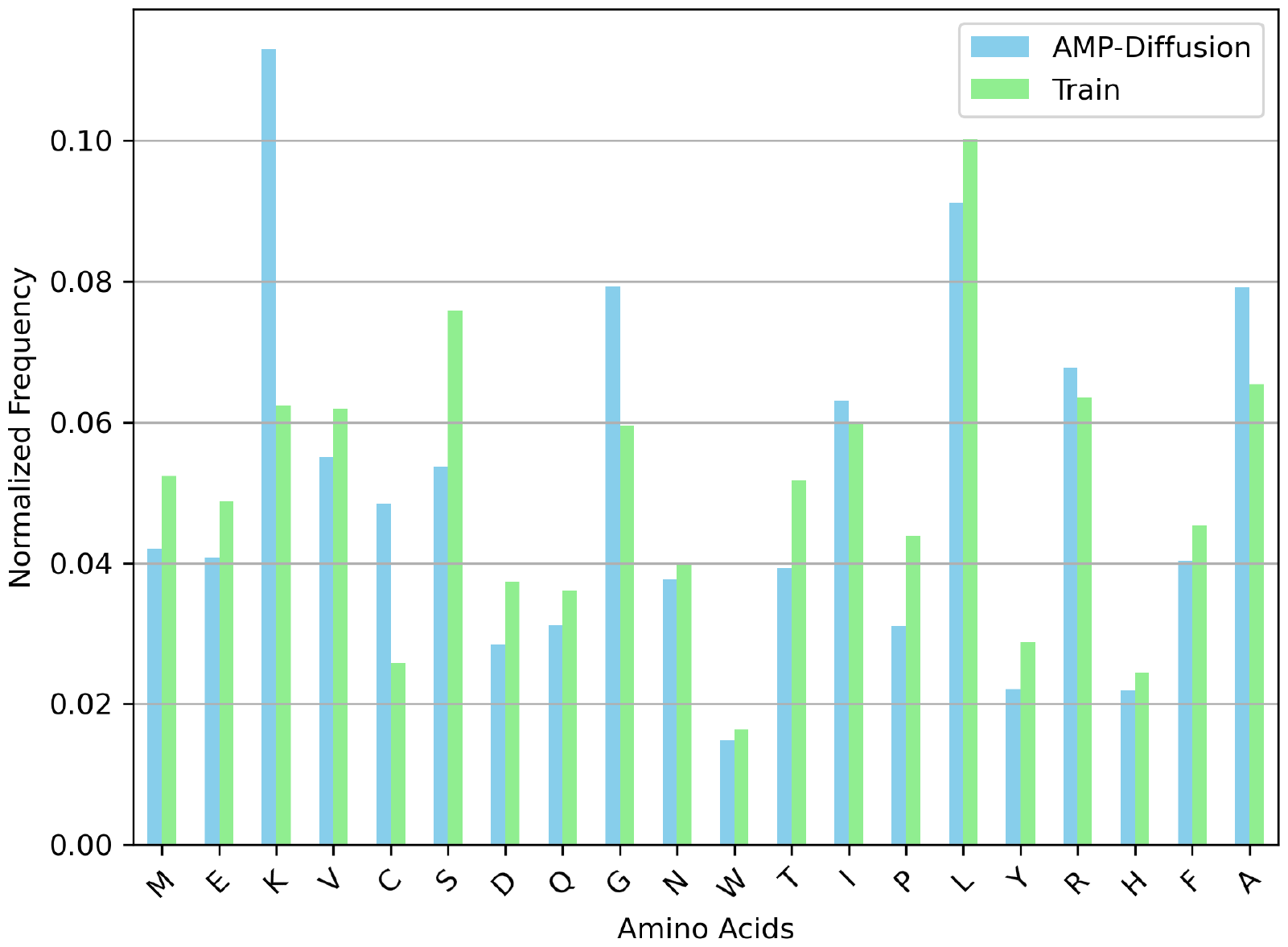
Normalized frequency comparison of standard amino acids between training data (green) and AMP-Diffusion generated peptides (blue).

**Figure 5:**
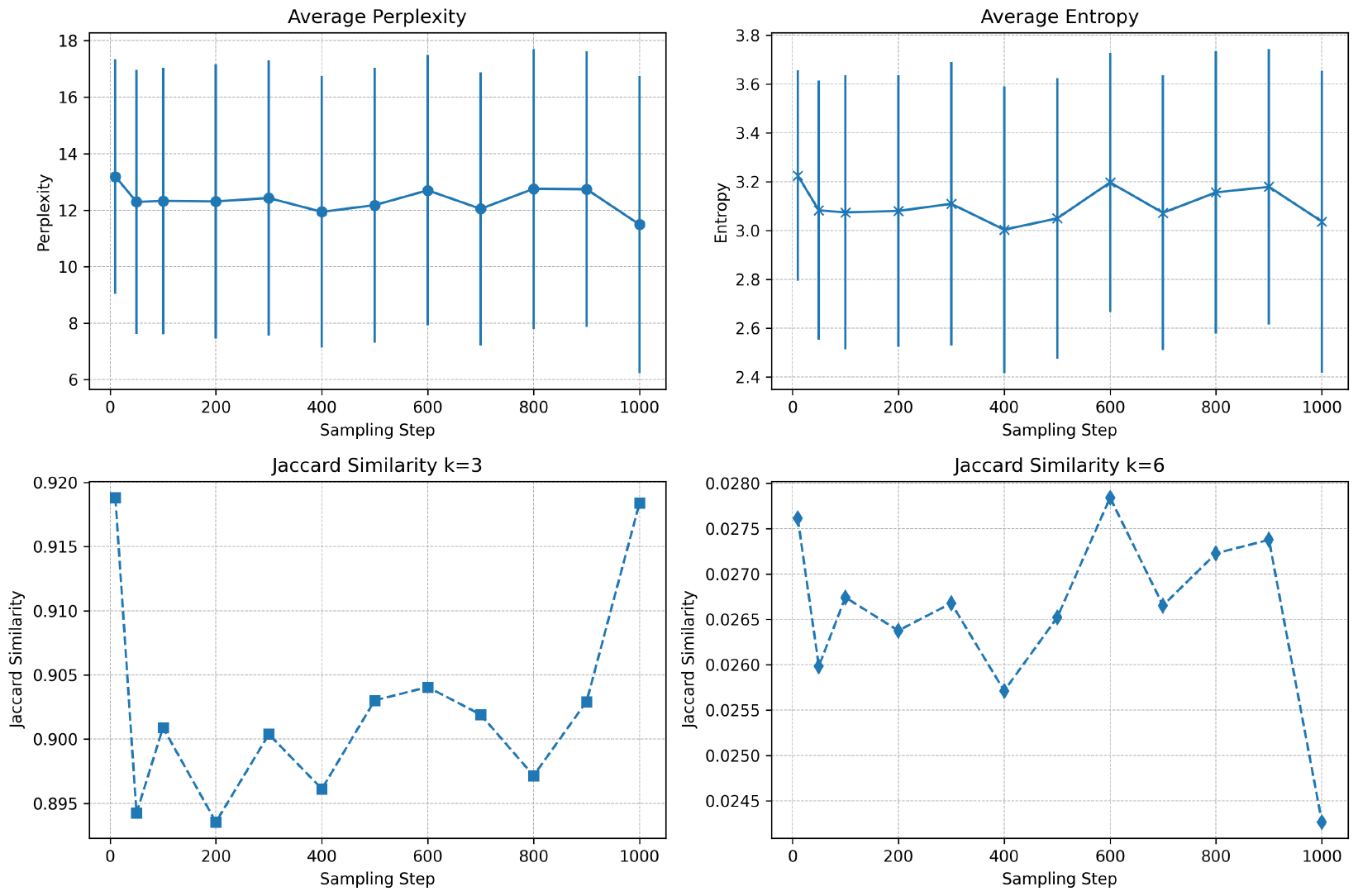
Evaluation of perplexity and diversity metrics across various sampling steps of the DDIM sampler. Six distinct sampling steps were examined, and the results indicate minimal variation in the assessed metrics across these steps.

## 4 Conclusion

This study presents an innovative approach to AMP design, combining the strengths of a pLM with a diffusion model. The pre-trained knowledge embedded within the pLM significantly enhances the diffusion model’s capacity to capture the essence of AMPs within the latent space. The peptides generated through this hybrid model demonstrate a statistically robust performance across multiple evaluation metrics and exhibit physicochemical properties that not only mirror those of real AMPs but also show potential improvements. The versatility of the framework is notable, with potential applications extending to general protein design tasks. Future research directions include experimental validation of the antimicrobial activities of the generated peptides, further exploration of the model’s ability to tailor peptides with specific desired properties, and broadening the use of our model architecture to other areas of protein engineering. Finally, future exploration of model parameters and the inclusion of additional conditioning data present opportunities to fine-tune the model for optimal performance and enhanced sequence diversity. Overall, AMP-Diffusion represents a successful integration of pLMs with diffusion models, offering a promising avenue for future innovation in protein design and laying the groundwork for future enhancements and applications in this ever-growing field of research.

## Acknowledgement

The authors extend their gratitude to the Duke Compute Cluster, Oracle Cloud, and Mark III Systems for their generous computational support, which was pivotal for this project. Heartfelt thanks are also given to the anonymous reviewers from ISMB 2023 and the NeurIPS 2023 Generative AI and Biology Workshop, whose insightful feedback and constructive suggestions significantly enhanced the quality of this work. Lastly, the team acknowledges the collective support and enriching discussions with the members of the Programmable Biology Group at Duke University.

## 5 Supplementary Material

### 5.1 Implementation Details

AMP-Diffusion was trained on four Nvidia A10 GPUs with batch size of 64 and learning rate 9.93 *×* 10^*−*4^. The Adam optimizer (28) was used for optimization. Hyperparameter tuning, specifically for the learning rate and batch size, was conducted over 100 runs, leveraging the Bayesian optimization method facilitated by Wandb (29). The training regimen incorporated a cosine learning rate schedule and utilized the *l*2 loss for the denoising component.

For the generation phase, a total of 50,000 samples were produced across 500 rounds, with each iteration yielding 100 samples. The length of these samples was randomly determined, ranging between 15 to 40 amino acids. In cases where the amino acid X appeared, which the HydrAMP classifier cannot process, it was replaced with a random standard amino acid for evaluation purposes. For other benchmarking models, we directly used sequences generated by the HydrAMP (19) team. The entire implementation was executed using PyTorch 2.01 and Python 3.10.10.

### 5.2 Related Works

Traditional computational methodologies primarily utilize classifiers to ascertain the AMP status of specific peptide sequences or to forecast properties such as cytotoxicity or peptide activity (30; 17). While these classifier-centric techniques are instrumental in segmenting peptides based on their antimicrobial traits, their scope is confined to predicting extant attributes of input peptides, rendering them inept at generating *de novo* AMP sequences. In a parallel vein, quantitative structure-activity relationship (QSAR) models endeavor to prognosticate the AMP status of peptide sequences (31; 32). These models operate by pinpointing structural features indicative of AMP status and subsequently “scoring” peptides based on these features. However, akin to classifier models, QSAR models are constrained to existing peptides, curtailing their utility in spawning novel AMPs.

In the pursuit of novel AMP synthesis, generative models have emerged as a promising avenue. Generative adversarial networks (GANs) (33) have been explored for their potential in AMP generation. These networks, particularly the bidirectional conditional GANs (BiCGANs), have been adapted to the unique challenges of AMP design. Oort et al. demonstrated the utility of BiCGANs in mapping AMP sequences into the generator’s latent space, thereby enabling intricate operations tailored to AMP characteristics, such as landmark creation and specific sequence manipulations (21). However, the inherent complexities in training GANs and their limited diversity in generation have been noted as challenges in the context of AMP design.

Variational autoencoders (VAEs), especially their conditional variants (cVAEs), have also been at the forefront of AMP-centric generative research. While vanilla VAEs offer a probabilistic approach to mapping input AMP data to latent spaces, cVAEs introduce conditional information, enhancing the specificity and control in AMP generation. Several studies have showcased the prowess of modified cVAEs in generating AMPs, emphasizing their capability to produce peptides under specific conditions related to AMP properties (20; 19). While cVAEs are recognized for their diversity in AMP generation, challenges related to the fidelity of generated sequences persist.

### 5.3 Generation Embedding Visualization

Linear and nonlinear methods were employed to reduce the dimensionality of the embeddings for visualization purposes. The results from PCA and t-SNE are depicted in Figure 3, showing the embeddings from both the training data and AMPs generated by AMP-Diffusion within a two-dimensional ESM latent space. Both plots reveal a clear clustering and overlapping of the data points, reflecting a significant resemblance between the actual and generated AMPs. Additionally, cosine similarity was calculated between each generated embedding and its nearest neighbor in the training set, yielding an average value of *±* 0.098 0.048. These findings imply that the model may effectively capture and manifest the inherent characteristics of natural AMPs.

### 5.4 Amino Acid Frequency

At the single amino acid level, we showed the normalized frequency of standard amino acids across training and generation AMPs in Figure 4. It indicates a high level of congruence between the two datasets for most amino acids. Notably, the amino acid lysine (K) is an exception, exhibiting a significantly higher frequency in the generated AMPs compared to its natural occurrence in the training data. This observation suggests that while AMP-Diffusion largely adheres to the natural distribution patterns of amino acids, it has a propensity to incorporate lysine more frequently in its generation.

### 5.5 DDIM Sampling

One common challenge faced by DDPM is the prolonged sampling duration. To mitigate this, a plethora of samplers have been proposed, notable among them being the DDIM (14) and DPM-solver (34). In this study, we also incorporated the DDIM approach, which introduces a novel generative process by considering non-Markovian forward processes. This allows for the exploration of generative processes that can produce samples tailored to specific requirements by merely adjusting certain parameters. A salient feature of DDIM is its ability to accelerate the generative process by considering forward processes with lengths smaller than the total number of steps, leading to a significant boost in computational efficiency.

In our experiments, we evaluated the DDIM sampler across varying sampling steps [10, 50, 100, 200, 300, …, 1000], using perplexity and diversity as our primary metrics. For each time step, 10,000 samples are generated. Our findings indicate that the choice of sampling steps did not drastically affect Perplexity and Entropy in Figure 5. However, regarding Jaccard similarity of 3-mers and 6-mers, *k* = 600 might be the optimal configuration. Although the performance of sampling step 10 looks good, the model exhibited a propensity to generate sequences with recurring regions. While these preliminary results provide valuable insights, a more exhaustive examination of the generated sequences is warranted. Future research directions could also encompass the exploration of even faster sampling algorithms to further enhance the efficiency of DDPMs.

## Notes

### Competing Interest Statement

The authors have declared no competing interest.

